# Increasing testing throughput and case detection with a pooled-sample Bayesian approach in the context of COVID-19

**DOI:** 10.1101/2020.04.03.024216

**Authors:** Rodrigo Noriega, Matthew H. Samore

## Abstract

Rapid and widespread implementation of infectious disease surveillance is a critical component in the response to novel health threats. Molecular assays are the preferred method to detect a broad range of pathogens with high sensitivity and specificity. The implementation of molecular assay testing in a rapidly evolving public health emergency can be hindered by resource availability or technical constraints. In the context of the COVID-19 pandemic, the applicability of a pooled-sample testing protocol to screen large populations more rapidly and with limited resources is discussed. A Bayesian inference analysis in which hierarchical testing stages can have different sensitivities is implemented and benchmarked against early COVID-19 testing data. Optimal pool size and increases in throughput and case detection are calculated as a function of disease prevalence. Even for moderate losses in test sensitivity upon pooling, substantial increases in testing throughput and detection efficiency are predicted, suggesting that sample pooling is a viable avenue to circumvent current testing bottlenecks for COVID-19.

---

Emerging infectious diseases pose a global hazard to public health, as exemplified by the COVID-19 pandemic. Key epidemiologic strategies for control of community spread include contact tracing, case isolation, ring containment, and social distancing (*1–7*). The use of microbiological testing to identify disease cases is a crucially important element of these strategies. Some countries, including the US, experienced a shortage of kits needed for COVID-19 diagnosis, which resulted in the imposition of restrictive criteria to manage the selection of patients for testing. Constraints in the supply of kits had a particularly significant impact on testing of mildly symptomatic individuals, as well as asymptomatic contacts of confirmed cases. For some facilities that have been overwhelmed by demand for testing as the pandemic progressed, test throughput continues to be a limiting factor (*8–10*). Strategies for screening more individuals with a reduced burden on resources are highly desirable. Using a Bayesian formalism, a hierarchical testing protocol based on sample pooling is discussed. Anticipated benefits include easing the demand of constrained resources and enabling more efficient detection of a larger number of cases.

Molecular assays are the predominant testing method for viral and bacterial pathogens (*11–14*). Specifically, nucleic acid detection assays typically employ real-time polymerase chain reaction (RT-PCR) for DNA targets and reverse-transcription real-time PCR (rRT-PCT) for RNA targets (*15, 16*). The popularity of such testing platforms is due to **1)** their high sensitivity and specificity, **2)** the widespread access to sequencing and synthesis technologies for the identification of nucleic acid target sequences and probes, and **3)** the development of fast, user-friendly, and cost-effective equipment. While nucleic acid assays have powered a revolution in diagnostics and delivery of care for individual patients, their application in large-scale infectious disease surveillance is hampered partly by low throughput at a population level.

The information content of a diagnostic test can be evaluated with a Bayesian probability formalism in the context of an individual sample or for repeated sampling from the same patient (*17–19*) by taking into account the probability of detecting a positive case (assay sensitivity, or identification rate *P*_*id*_) and the probability of a positive result from healthy samples (false positive rate *P*_*fp*_). Bayesian inference requires the assessment of a “prior” probability to the presence of disease in a sample, *P*(*D*), which is updated to a “posterior” probability given a positive or negative test result with conditional probabilities *P*(*D*|+) or *P*(*D*|−) (**Eq. 1.a-b)**.

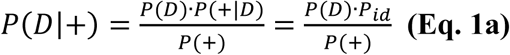

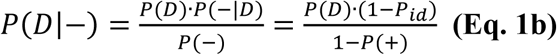

where *P*(+) and *P*(−) are the overall probabilities of the test yielding a positive or negative result, respectively. These tools can be extended to the somewhat counterintuitive situation where a diagnostic test is conducted on a sample pooled from multiple individuals. The motivation for sample pooling is to screen multiple patients simultaneously and reduce the burden on testing facilities working with limited resources. Pooling schemes have been developed since their introduction in the 1940s for syphilis testing, and have been applied to screen for and estimate prevalence rates of a variety of diseases (*20-24*). Here, a simple two-step hierarchical protocol first introduced by Dorfman is considered: *Samples from N*_*p*_ *patients are collected and randomly pooled into groups of n individual samples each. Pooled samples are interrogated with the diagnostic test. If pooled testing yields a negative result, no further testing is conducted. If pooled testing yields a positive result, all patients in that pool are tested individually.*

While statistical approaches have been focused on characterizing the performance of various pooling schemes, not all of them include non-ideal test parameters (*25*). Moreover, testing characteristics at the pooled- and individual-sample levels can be different. Here, the Bayesian inference approach in **Eq. 1.a-b** is modified to include differences in the assay sensitivity and overall probability of a positive result in pooled vs. individual tests (**Eq. S1-2**). Sensitivity loss is included as a reduction in the identification rate of pooled-sample tests by a scaling factor *γ*. Due to the exceptional specificities of nucleic acid assays, the false positive rate is assumed to remain unaffected. Importantly, the posterior probability assessed from a positive pooled test is used as a prior for follow-up individual tests, which yields additional information and enhances the Bayesian inference assessment of those cases (**Eq. S3**).

For a population of *N*_*p*_ patients divided into sample pools of size *n*, the average number of tests is the number of initial pooled tests plus the expected number of follow-up tests (**Eq. S4**). Throughput increase *χ* is expressed as the effective number of individuals screened by each diagnostic test (**Eq. S5**). The individual- and pooled-sample test characteristics determine the pool size that optimizes screening throughput as a function of average disease prevalence in the tested population.

The advantages of pooled-sample screening are discussed in the context of the rapidly evolving COVID-19 pandemic (caused by the novel coronavirus SARS-CoV-2) (*8, 26*). A recent rRT-PCR assay for COVID-19 reports an identification rate of 95% and no false positives after testing 310 samples including other respiratory pathogens (*27*). Given reported specificities of commercially-available respiratory panel assays (>99%), an estimated 1% false positive rate was included in the model results reported here. A moderate reduction in the identification rate for a pooled sample (*γ* = 0.9) was assumed – this variable is discussed in detail below. Consistent with similar implementations of Dorfman-type testing algorithms (*28*), substantial increases in testing throughput are predicted for low disease prevalence rates (*P*(*D*) ≤8%, **Fig. 1**), where throughput more than doubles and optimal pool sizes are 4 ≤ *n* ≤ 12. At intermediate prevalence rates 0.1 ≤ *P*(*D*) ≤ 0.2, the increase in throughput is moderate yet substantial (>30% increase in throughput) and pool sizes are small (*n* = 3). For high prevalence rates, pooling yields no improvement (optimal pool size *n* = 1). Average disease prevalence can be re-assessed as information is gained for the tested population to re-optimize pool size. Dynamic self-tuning is a feature of Bayesian inference, a significant asset when a close feedback loop is desirable.

**Fig. 1.**
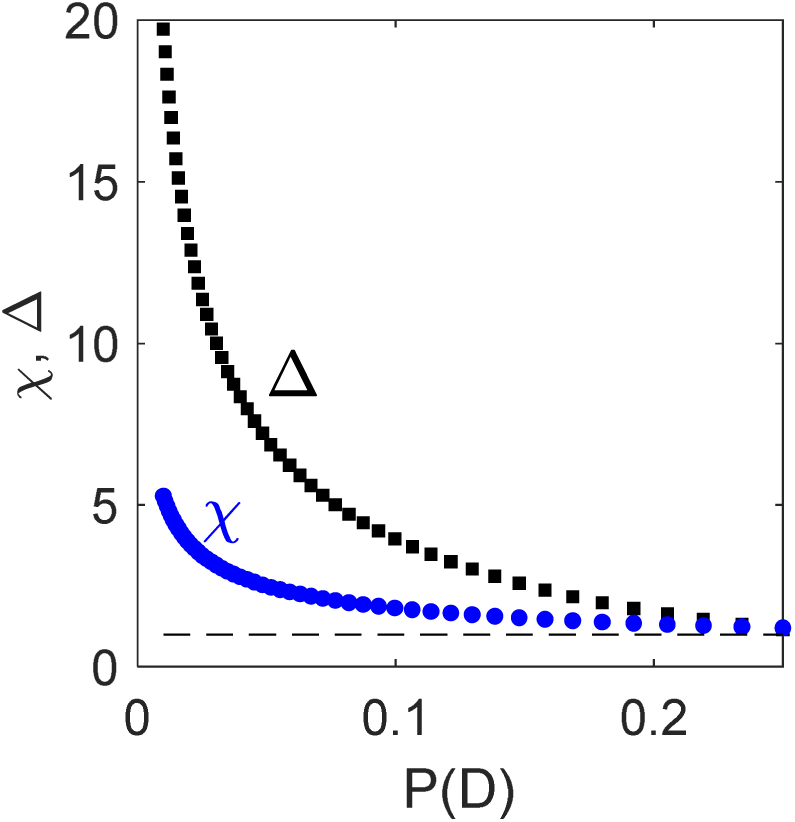
Number of patients screened by each diagnostic test (χ, blue circles) and the relative increase in detection-to-miss ratio for a pooled scheme (Δ, black squares), as a function of average disease prevalence. Gray line is the no-pooling scheme reference.

Besides testing throughput, it is informative to assess the increased ability to detect cases using a pooled-sample vs individual-sample scheme, which can be accomplished by comparing the ratio of detected to missed cases in each protocol. Importantly, resource constraints are incorporated into this comparison by accounting for unscreened cases in the standard 1:1 scheme (**Eq. S6-8**). The relative increase in the detection-to-miss ratio between the pooled and standard 1:1 schemes exhibits even more significant gains than those observed for testing throughput (**Fig. 1**).

Loss in screening power associated with sample pooling is assessed by **1)** estimating the portion of cases that would have been identified in an individual test but missed by pooled screening, and **2)** determining the information gained for patients whose pooled screening result was negative.

A reduction in overall pathogen concentration due to pooling in conditions of low disease prevalence can decrease the test’s identification rate, although it is manageable when targeting infectious diseases for which typical pathogen concentrations are non-negligible (*23*). This concern is examined with a set of 186 positive rRT-PCR diagnostic test results for COVID-19 (**Fig. 2**), which include nasopharyngeal swab, oropharyngeal swab, and bronchoalveolar lavage (*29*). Tests have a mean cycle threshold value of ⟨*C*_*t*_⟩ = 24.6, and a positive test result is defined as a rRT-PCR reaction with *C*_*t*_ ≤ 42 – in agreement with early reports for nasal swab COVID-19 rRT-PCR tests with ⟨*C*_*t*_⟩ = 24.3 and viral loads of 1.4 × 10^6^ copies/mL (*30*). From these data, the portion of samples that would have had a positive test result even if pooled with an entirely healthy population is estimated using a pool size *n* = 12 and rRT-PCR geometric efficiencies of ϵ = 1.7 − 2 per cycle. A pooled screening protocol would have detected 95.7-96.8% of these cases (178-180 out of 186). Further support for moderate sensitivity loss can be achieved by dividing the distribution of *C*_*t*_ values for the test-positive samples into three subpopulations. For the same pooling and rRT-PCR efficiencies stated previously, >95% of the broadest, lowest-load population would be detected.

**Fig. 2.**
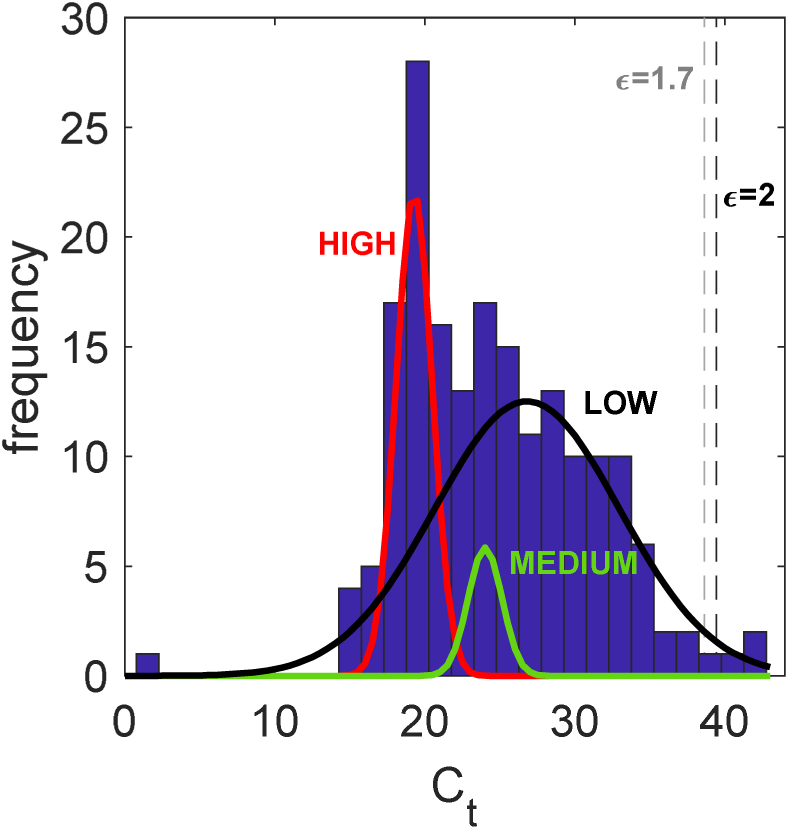
Cycle threshold values for rRT-PCR reactions for confirmed COVID-19 cases (*27*), described by subpopulations with low, medium, and high viral load. Sensitivity cutoffs shown as vertical lines.

The main benefit of hierarchical pooled-sample testing protocols is the ability to screen a larger portion of the population and detect more positive cases. The relative increase in case detection is given by the ratio of the number of cases detected in a pooled setting and the number of cases detected if the same number of tests were used on single patients – the multiplicative factor in testing throughput scaled by the loss in sensitivity, *χ* · *γ*. Loss in sensitivity also leads to more screened-yet-missed cases – even in the absence of pooling, diagnostic tests yield false negatives and increasing those odds should not be considered lightly. In the range of disease prevalence with moderate to high increase in throughput and compared to individual testing of every patient, a *γ* = 0.9 sensitivity loss leads to a relatively low increase in posterior probability of disease after a negative test outcome, *P*(*D*|−) (**Fig. 3**). While sensitivity losses decrease the ability to confidently screen healthy individuals, the threshold for this tradeoff depends on the situation where the screening protocol is deployed. Effective risk communication to screened individuals is needed to prevent an outsized sense of security after a negative pooled-test result – e.g., vigilance to symptom development triggers individual testing.

**Fig. 3.**
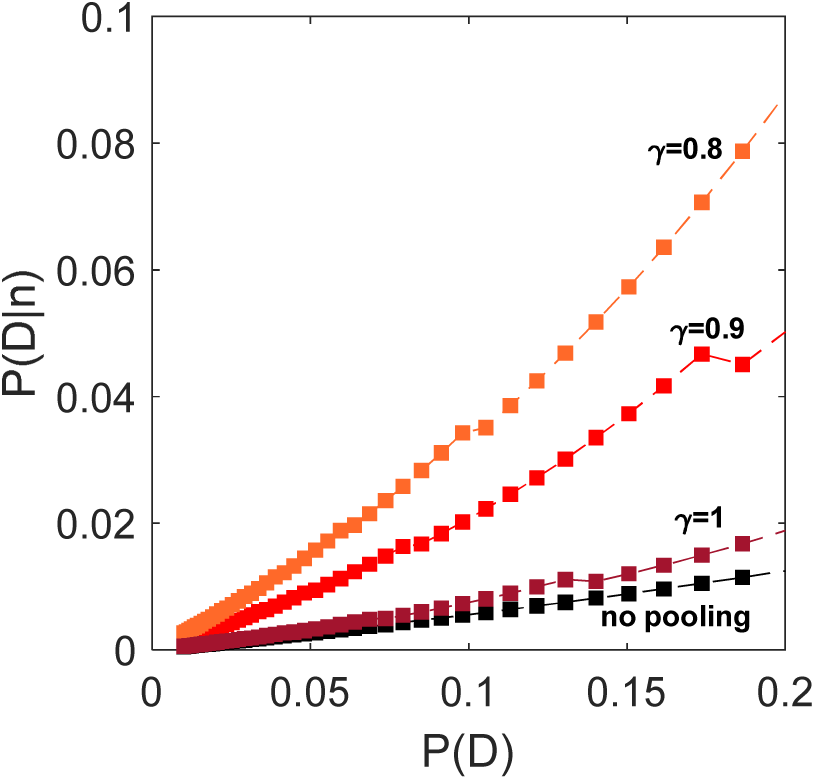
Probability of an individual being infected even though their pooled-sample test gave a negative result, as a function of background disease prevalence and for different values of sensitivity loss.

This protocol is amenable to HIPAA regulations – in fact, pooling has been implemented by state laboratories in the recent past (*31*)– and requires limited additional sample processing. Refining the dependence of sensitivity loss with pool size and coupling to modeling strategies that inform population sampling and prior probabilities will provide further screening improvements. From an epidemiological surveillance standpoint, increased detection of positive cases in a larger portion of the population denotes that a greater fraction of infectious individuals can be isolated. However, pooled testing in less useful in an inpatient clinical setting where the highest sensitivity is needed to minimize risk of hospital transmission from non-isolated patients. As with any change in the delivery of medical care, a discussion including community stakeholders is paramount.

In summary, a pooled testing strategy has the potential to enhance comprehensive surveillance of SARS-CoV-2 particularly when test kits are in short supply. The benefits of surveillance are greatest in the early phases of community spread. Thus, improving the capacity for high-throughput testing has the highest impact when prevalence is low enough that pooled sampling is most beneficial. The ratio of confirmed COVID-19 cases to tests performed varies by country, but it appears that aggressive testing strategies yield a low enough prevalence to benefit from pooled-sample screening – e.g., South Korea’s is 3% (8.3k/270k as of 3/16/2020) (*32*). While the development of clinical prediction rules and non-testing screening are critical to any epidemiological response, dealing with a novel disease for which data is still sparse and testing capabilities are limited means that maximizing the impact of each individual test can benefit the continued refinement of our strategy.

## Acknowledgments

Valuable discussions with Dr. Kimberly Hanson and Dr. Lindsay T. Keegan.

## Author contributions

R.N. conceived the idea, performed the analysis, prepared the figures, and drafted the manuscript. R.N. and M.H.S. developed the idea and edited the manuscript;

## Data and materials availability

All data is available in the main text or the supplementary materials.

## Materials and Methods

### Bayesian inference implementation

The process for assessing the posterior probabilities for individuals whose pooled test yielded a positive or negative result (**Eq. S1.a-b**)

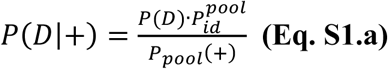

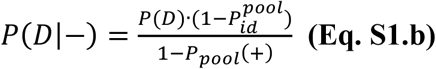

includes the a reduction in the identification rate of the pooled-sample diagnostic test by a factor *γ* compared to an individual test, so that 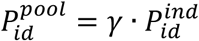 One must also consider the probability of a positive result in a pooled sample. Assuming that every individual has a prior probability of being infected equal to some background average *P*(*D*), the probability of a positive test result in a pool of size *n* is the probability of having a nonzero number of positive individual samples times the pooled-test identification rate plus the probability of a completely-healthy sample pool yielding a false positive (**Eq. S2**):

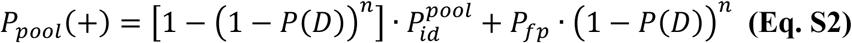

The posterior probability *P*(*D*|+) from the pooled test with a positive outcome can be used as a prior for the follow-up individual test Bayesian inference (**Eq. S3.a-b**).

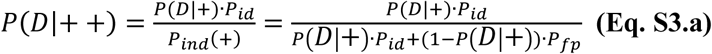

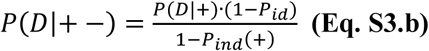

where the overall probability of an individual’s test being positive includes the probability of correctly identifying a positive sample *P*(*D*|+) · *P*_*id*_ plus the probability of a negative sample yielding a false positive (1 − *P*(*D*|+)) · *P*_*fp*_.

For a population of *N*_*p*_ patients divided into sample pools of size *n*, the average number of tests ⟨*N*_*test*_⟩ is given by the number of initial pooled tests plus the expected number of follow-up tests (**Eq. S4**):

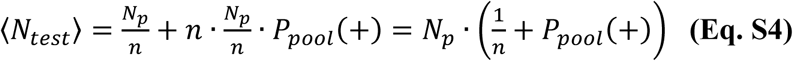

The throughput increase is expressed as a multiplicative factor representing how many individuals were screened by the use of each diagnostic test (**Eq. S5**).

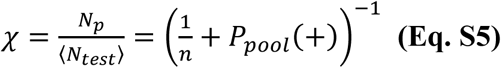

#### Full vs. partial optimization

In a fully optimized scheme a pooled-sample’s positive result triggers a cascade of smaller-sized pools, minimizing the number of tests performed. However, an intermediate improvement is chosen due to possible adverse outcomes of a lengthier process with several pooling and Bayesian inference steps – e.g., possible delay of necessary care, increased exposure to infected individuals.

### Advantages in testing throughput and case detection

We can estimate the ratio of detected-to-missed (including unscreened) cases for each protocol with the following simplified analysis:

<u>**In the pooled scheme**</u>, the number of detected cases are the total number of screened individuals with the disease ⟨*N*_*test*_⟩*χP*(*D*) multiplied by the effective detection probability *γ* · 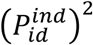, which includes the sequential detection probability at the pooled stage and the follow-up individual stage. The number of missed cases would thus be the difference between detected and total cases, 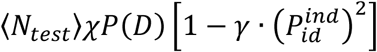 The number of false positives is the number of individuals in all-healthy pools yielding a false positive and whose individual test is also a false positive – 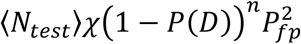 – plus the healthy individuals pooled with a diseased sample that is detected at the pooled stage and whose follow up test is a false positive, 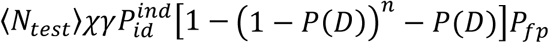 The detection-to-miss ratio *θ*_*pool*_ can be expressed as

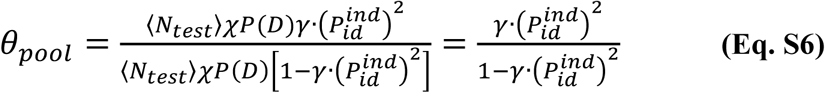

<u>**In a standard 1:1 scheme**</u>, the number of detected cases are the number of tested individuals with the disease ⟨*N*_*test*_⟩*P*(*D*) times the detection efficiency of the individual test 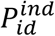 The number of missed cases is thus 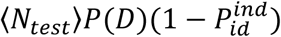 The number of false positives is ⟨*N*_*test*_⟩[1 − *P*(*D*)]*P*_*fp*_. The number of unscreened cases that carry the disease is ⟨*N*_*test*_⟩(*χ* − 1) *P*(*D*). The detection-to-miss ratio – considering the population sampled in the pooled scheme – is given by

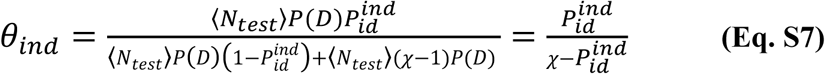

The relative increase in the detection-to-miss ratio between the pooled and standard 1:1 schemes is thus given by

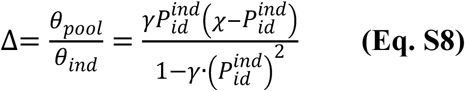

To further illustrate this, consider a disease including asymptomatic-yet-infectious individuals and for which clinical predictions are at an early stage, preventing effective triage against conditions presenting similar symptoms. A screening point must decide how to use limited resources (e.g., 4,000 available tests) to detect the maximum number of cases in a population at 5% risk. Using the test characteristics described in the main text 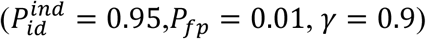, it is possible to compare the pooled-sample screening protocol vs. standard 1:1: testing. On average, a pooled-sample approach allows testing 10,000 individuals, detecting 406 cases while missing 94 and yielding 18 false positives; conversely, a standard 1:1 approach would test 4,000 individuals and detect 190 cases, miss 10, yield 38 false positives, and leave 300 positive cases untested and thus undetected.

What if the rRT-PCR test sensitivity is much lower than expected – say 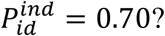 Let us also consider a much more pessimistic estimate for sensitivity loss, *γ* = 0.8 and no change in the false positive rate (*P*_*fp*_ = 0.01). For the same 5% prevalence in the population, the screening point with access to 4,000 tests would screen 12,550 individuals (627 of which are infected) using a pooling scheme (for these new parameters, optimal pool size is *n* = 7). It would detect 246 and miss 381. A 1:1 testing scheme would identify 140 cases, miss 60, and leave 427 untested. Not surprisingly, the number of false negatives increases substantially <u>in both scenarios</u> due to the lower starting point for the test’s sensitivity. However, in a pooled scheme 381 infected individuals are still at risk of spreading the disease in the community, while in the 1:1 scheme 487 infected individuals remain at risk of further community spread. As mentioned in the main text, effective risk communication is a critical component of any large-scale screening effort with imperfect tests. Symptomatic individuals should be considered at increased risk even after a negative test result, and other diagnostic avenues could be used (e.g., chest CT).

